# Catalog of gut microbiota alterations associated with anticancer therapies across multiple cancer types

**DOI:** 10.64898/2026.05.08.723707

**Authors:** Kyoko Kurihara, Shunsuke A. Sakai, Kentaro Sawada, Naoko Iida, Satoshi Horasawa, Takao Fujisawa, Yoshiaki Nakamura, Shun-Ichiro Kageyama, Hideaki Bando, Takayuki Yoshino, Katsuya Tsuchihara, Riu Yamashita

## Abstract

**Background:** Anticancer therapies can alter the gut microbiota and may affect gut bacteria associated with treatment response. However, most longitudinal studies have focused on specific cancer types or individual treatment regimens, and systematic analyses across diverse cancer therapies remain limited. We analyzed longitudinal fecal microbiota profiles using 16S ribosomal RNA gene amplicon sequencing in the pan-cancer SCRUM-Japan MONSTAR-SCREEN cohort. We included 528 paired pre- and post-treatment fecal samples from 264 patients with advanced solid tumors across 18 cancer types and characterized the gut microbiota alterations associated with 22 anticancer drugs and related clinical factors.

**Results:** Across the cohort, Shannon diversity did not significantly change after treatment (mean, 3.81 vs. 3.78; *P* = 0.58), and pre- and post-treatment samples exhibited no clear separation in ordination space. However, within-patient analysis detected a subtle but significant longitudinal microbiota shift (paired PERMANOVA, *P* = 0.0001), highlighting the importance of accounting for paired sampling. Clustering of genus-level compositional alterations revealed patient groups with distinct degrees of microbiota alteration, with the largest shifts associated with antibiotic exposure, transition from normal stool to diarrhea, and specific treatment regimens. Multivariable regression analysis of 22 anticancer drugs identified drug–bacteria associations and demonstrated that drugs with similar mechanisms of action, including epidermal growth factor receptor (EGFR) inhibitors and immune checkpoint inhibitors, exhibited similar microbiota change profiles. Targeted analyses highlighted concordant reductions in the Christensenellaceae R-7 group among EGFR inhibitor-exposed patients and depletion of *Faecalibacterium* among immune checkpoint inhibitor-exposed patients.

**Conclusions:** This study provides a cross-cancer catalog of microbiota alterations associated with anticancer therapies and highlights therapy-related shifts in the gut ecosystem, including patterns shared by drugs with similar mechanisms of action.

## Background

The gut microbiota has emerged as a crucial modifier of cancer treatment outcomes [1]. Pre-treatment microbial composition has been associated with immune checkpoint inhibitor (ICI) efficacy across multiple cancer types [2–6]. Fecal microbiota transplantation from responding donors has induced clinical responses in a subset of patients with melanoma refractory to anti–programmed cell death protein 1 (anti–PD-1) therapy [7, 8], supporting the therapeutic relevance of microbiome modulation. Microbiota-mediated effects may extend beyond immunotherapy because commensal bacteria and microbial metabolites can modulate the response to platinum chemotherapy [9, 10]. Emerging clinical evidence has further associated gut microbial features with the outcomes of targeted therapies, including poly(ADP-ribose) polymerase inhibitor maintenance therapy in ovarian cancer [11].

As microbiome-directed strategies advance toward clinical application, it is increasingly crucial to determine when and in which treatment contexts these interventions may be most effective. Cancer therapies may affect the gut ecosystem through multiple mechanisms, including direct effects on bacterial growth [12], microbial transformation of therapeutic compounds that may create selective pressure on bacterial communities [13–15], and treatment-related gastrointestinal toxicities, such as mucositis and diarrhea [16]. A longitudinal understanding of therapy-associated microbial dynamics is needed to define rational targets and timing for future microbiome-based interventions.

However, comprehensive studies assessing gut microbiota alterations across diverse cancer therapies within a unified analytical framework remain limited. Existing studies have largely focused on individual cancer types or specific treatment settings, such as ICI-treated melanoma, endocrine therapy in breast cancer, ICI-associated resistance or immune-related adverse events, and capecitabine-treated colorectal cancer [17–20]. Methodological heterogeneity across studies, including differences in fecal sample preservation, DNA extraction protocols, and 16S ribosomal RNA (rRNA) gene target regions, can confound cross-study comparisons [21–23]. Therefore, a standardized longitudinal analysis across multiple malignancies and therapeutic agents is needed to obtain a more coherent view of treatment-associated gut microbiota alterations.

In this study, we used clinical and longitudinal fecal microbiota data from the SCRUM-Japan MONSTAR-SCREEN cohort [24, 25] to systematically characterize the gut microbiota alterations associated with anticancer treatment. We analyzed paired pre- and post-treatment fecal samples from 264 patients with 18 cancer types and used multivariable regression models to assess the association between alterations in bacterial genera and exposure to 22 anticancer drugs. This analysis generated a cross-cancer catalog of drug–bacteria associations, providing a systematic reference for therapy-associated microbiota alterations.

## Methods

### Study design and cohort

This study was based on the SCRUM-Japan MONSTAR-SCREEN project [24, 25], a nationwide multicenter observational study of advanced solid tumors, excluding lung cancer. A total of 2,196 patients were enrolled from 31 institutions in Japan between October 2019 and September 2021. Clinical data, questionnaire-based lifestyle data, tumor tissue DNA, plasma circulating tumor DNA, and fecal samples were collected.

The primary eligibility criteria for MONSTAR-SCREEN were as follows: histopathologically confirmed unresectable or metastatic solid tumors; receipt or planned receipt of systemic therapy; age ≥ 16 years; Eastern Cooperative Oncology Group performance status of 0 or 1; adequate organ function; and life expectancy of at least 12 weeks. All the participants provided written informed consent. This study adhered to the Declaration of Helsinki and the Japanese Ethical Guidelines for Medical and Health Research Involving Human Subjects. The study protocol was approved by the institutional review board at each participating site and registered in the University Hospital Medical Information Network Clinical Trials Registry (UMIN000036749).

### Clinical data collection

We retrieved clinicopathological data from the electronic data capture system and used data available as of October 31, 2023. The dataset included age, sex, body mass index (BMI), cancer type, primary tumor status, treatment line details (start date, disease progression date, treatment discontinuation date, and discontinuation reason), treatment regimen, antibiotic exposure, and fecal sampling data (collection date and Bristol stool scale). For patients treated with ICIs, data on proton pump inhibitor (PPI) use was also collected.

### Fecal sample collection, DNA extraction, and 16S rRNA gene sequencing

Fecal samples were processed as previously described [24]. Briefly, fecal samples were collected at each participating institution using a commercial sampling kit containing a preservative solution (TechnoSuruga Laboratory Co., Ltd.) and transported to a central laboratory for 16S rRNA gene amplicon sequencing. Fecal samples were stored in a preservative solution at room temperature for up to 7 days and subsequently stored at -80 °C until DNA extraction. DNA was extracted in a central laboratory, following the manufacturer’s protocol. Briefly, 150 μL of stool suspension was processed using the NucleoSpin 96 Soil kit (Macherey-Nagel). DNA was eluted in 100 μL, purified using Agencourt AMPure XP beads (Beckman Coulter), and quantified using the PicoGreen dsDNA Assay Kit (Thermo Fisher Scientific).

For 16S rRNA gene amplicon sequencing, 1 ng of purified DNA was used as input for the first-round polymerase chain reaction (PCR) targeting the V3–V4 region using the 16S Metagenomic Library Construction Kit (Takara Bio). Amplicons were purified using Agencourt AMPure XP beads (Beckman Coulter), followed by index PCR using the Nextera XT Index Kit (Illumina). Indexed libraries were purified using Agencourt AMPure XP beads (Beckman Coulter) and quantified using the Quant-iT PicoGreen dsDNA Assay Kit (Thermo Fisher Scientific). Equimolar amounts of the indexed PCR products were pooled to generate the final sequencing library. The library size and concentration were assessed using a TapeStation system (Agilent Technologies). Sequencing was performed on the MiSeq platform (Illumina) with 250-bp paired-end reads using the MiSeq Reagent Kit v3 (Illumina), with approximately 40–50 % PhiX spike-in (Illumina).

### Processing of 16S rRNA gene sequence data

Paired-end FASTQ files were processed using QIIME 2 (version 2022.2) [26]. Reads were denoised with DADA2 to generate amplicon sequence variants (ASVs), with 22 bases trimmed from the 5′ end of both forward and reverse reads and truncation at 250 and 240 bases, respectively. Taxonomy was assigned using a pre-trained SILVA 138 99% Naive Bayes classifier implemented in the q2-feature-classifier. Samples with < 1,000 total reads were excluded. Genus-and ASV-level feature tables were exported for downstream analyses. For analyses requiring an even sequencing depth, samples were rarefied to 38,807 reads, corresponding to the minimum sequencing depth across the retained samples.

### Selection of patients with paired pre- and post-treatment fecal samples

Among the 2,196 patients enrolled in MONSTAR-SCREEN, we selected those with both a pre-treatment fecal sample collected within 14 days before or on the day of treatment initiation and a post-treatment fecal sample collected on or within 28 days after confirmed disease progression or treatment discontinuation. For treatment lines discontinued because of disease progression, the confirmed progression date was used as the reference date; for all other treatment lines, the treatment discontinuation date was used. When multiple treatment lines were available for the same patient, the later treatment lines were excluded. Ultimately, 264 patients with paired samples (n = 528) were included in the analysis.

### Alpha and beta diversity analyses

Alpha diversity was assessed using the Shannon index [27] calculated from the rarefied ASV count table with scikit-bio (version 0.6.2). Differences in the Shannon index between paired pre- and post-treatment samples were assessed using the Wilcoxon signed-rank test implemented in SciPy (version 1.15.3).

For beta diversity analysis, the Aitchison distance was calculated from the non-rarefied genus-level count table. Genera with prevalence < 10 % were excluded, leaving 181 genera for analysis. After adding a pseudocount of 1, the counts were converted to relative abundances and subjected to centered log-ratio (CLR) transformation [28]. Euclidean distances in the CLR-transformed space were used as Aitchison distances. Principal coordinate analysis (PCoA) was performed for ordination. Differences between pre- and post-treatment samples were tested primarily using paired permutational multivariate analysis of variance (PERMANOVA) in R (vegan version 2.6-10), with 99,999 permutations constrained within subjects using permute (version 0.9-7). Unpaired PERMANOVA was performed in Python using scikit-bio with 999 permutations for comparison. To visualize the distribution of within-patient Aitchison distances, a two-component Gaussian mixture model was fitted using scikit-learn (version 1.5.1).

### Unsupervised clustering and association with clinical features

For the clustering analysis, we calculated the within-patient change in CLR-transformed abundance for each genus between the pre- and post-treatment samples. After normalization, a paired-difference matrix was used for unsupervised patient clustering using ComplexHeatmap (version 2.22.0), applying six k-means clusters (column_km = 6) and 100 repeats (column_km_repeats = 100). The column ordering within the heatmap was based on Ward’s minimum variance method (clustering_method_columns = “ward.D2”). Differences in the Aitchison distance across clusters were assessed using the Kruskal–Wallis test in SciPy.

For each cluster, bacterial alterations were assessed using ALDEx2 (version 1.22.0) [30] with a filtered non-rarefied genus-level count table, excluding genera with a prevalence of < 10 %. Paired differential abundance analysis was conducted using the following parameters—mc.samples = 128, test = “t”, effect = TRUE, denom = “all”, paired.test = TRUE, and gamma = NULL. Genera with a Benjamini–Hochberg-adjusted *P* value (we.eBH) < 0.05 were considered significant.

The associations between clusters and clinical characteristics, including sex, age, BMI, cancer type, Bristol stool scale category, presence of the primary tumor, antibiotic exposure, and anticancer drug use, were assessed by comparing each cluster with all the remaining clusters combined. Bristol stool scale scores of 1–5 and 6–7 were classified as normal and diarrheal, respectively. Transitions between pre- and post-treatment samples were categorized as normal-to-normal, normal-to-diarrhea, diarrhea-to-normal, or diarrhea-to-diarrhea. Antibiotic exposure was grouped as presented in Table S2. Continuous variables were analyzed using the Mann–Whitney U test, with effect sizes defined as the differences in medians. Categorical variables were analyzed using Fisher’s exact test, with odds ratios reported as effect sizes. Categories represented by fewer than five patients were excluded, and for variables with over two categories, each remaining category was tested against all others. Only categorical variables with *P* < 0.05 were visualized.

### Multivariable regression analysis

Separate multivariable linear regression models were fitted for each of the 66 genera with a pre-treatment prevalence of at least 50 %, using the within-patient change in CLR-transformed abundance (post-treatment minus pre-treatment) as the outcome. For each genus, a single model simultaneously included 22 anticancer drugs administered to at least five patients. Covariates were selected based on the preceding cluster association analysis, retaining variables with *P* < 0.05. Cancer type was excluded because of multicollinearity, and all antibiotic classes used in at least five patients were included. The final models included sex, age, use of cephem, new quinolone, penicillin, tetracycline, sulfamethoxazole/trimethoprim, and the transition from normal stool to diarrhea. Three patients with missing antibiotic data were excluded. Multicollinearity was assessed using variance inflation factors (VIF) (Table S4). Models were fitted in statsmodels (version 0.14.5) with heteroskedasticity-robust standard errors (cov_type = “HC3”) as follows:

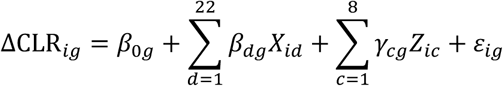

where ΔCLR*_ig_* denotes the within-patient change in CLR-transformed abundance of genus *g* for patient *i*, *X*_*id*_ denotes exposure to anticancer drug *d*, *Z*_*ic*_ denotes covariate *c*, and ε_*ig*_ denotes the residual error term. Nominal *P* values were reported. The estimated regression coefficients were assembled into a drug-by-genus coefficient matrix to summarize the drug–bacteria associations. Dimensionality reduction of the coefficient matrix was performed using uniform manifold approximation and projection (UMAP) (umap-learn version 0.5.7; n_neighbors = 5, min_dist = 0.5).

### Data visualization

Data visualization was performed using matplotlib (version 3.10.3) and seaborn (version 0.13.2) in Python 3.12.4, and ggplot2 (version 3.5.2) and ComplexHeatmap (version 2.22.0) in R 4.4.1.

### Use of AI tools

ChatGPT (OpenAI) was used to assist with the translation, manuscript language editing, and code editing. All AI-assisted outputs were reviewed and revised by the authors, who took full responsibility for the final manuscript and code.

## Results

### Heterogeneous longitudinal shifts in the gut microbiome across the cohort

We analyzed the MONSTAR-SCREEN cohort to assess the longitudinal effects of anticancer therapies on the gut microbiome (n = 264 patients with paired pre- and post-treatment fecal samples; 528 samples in total; Figure 1A and 1B). The cohort was heterogeneous, spanning 18 cancer types, with pancreatic (n = 59) and colorectal cancers (n = 50) being the most prevalent (Table 1; Figure S1A and Table S1). Across the cohort, pre- and post-treatment fecal microbiome profiles did not exhibit clear separation. No significant change in alpha diversity, as measured by the Shannon index, was observed across the full cohort (mean, 3.81 vs. 3.78; *P* = 0.58; Figure 1C, left).

**Figure 1.**
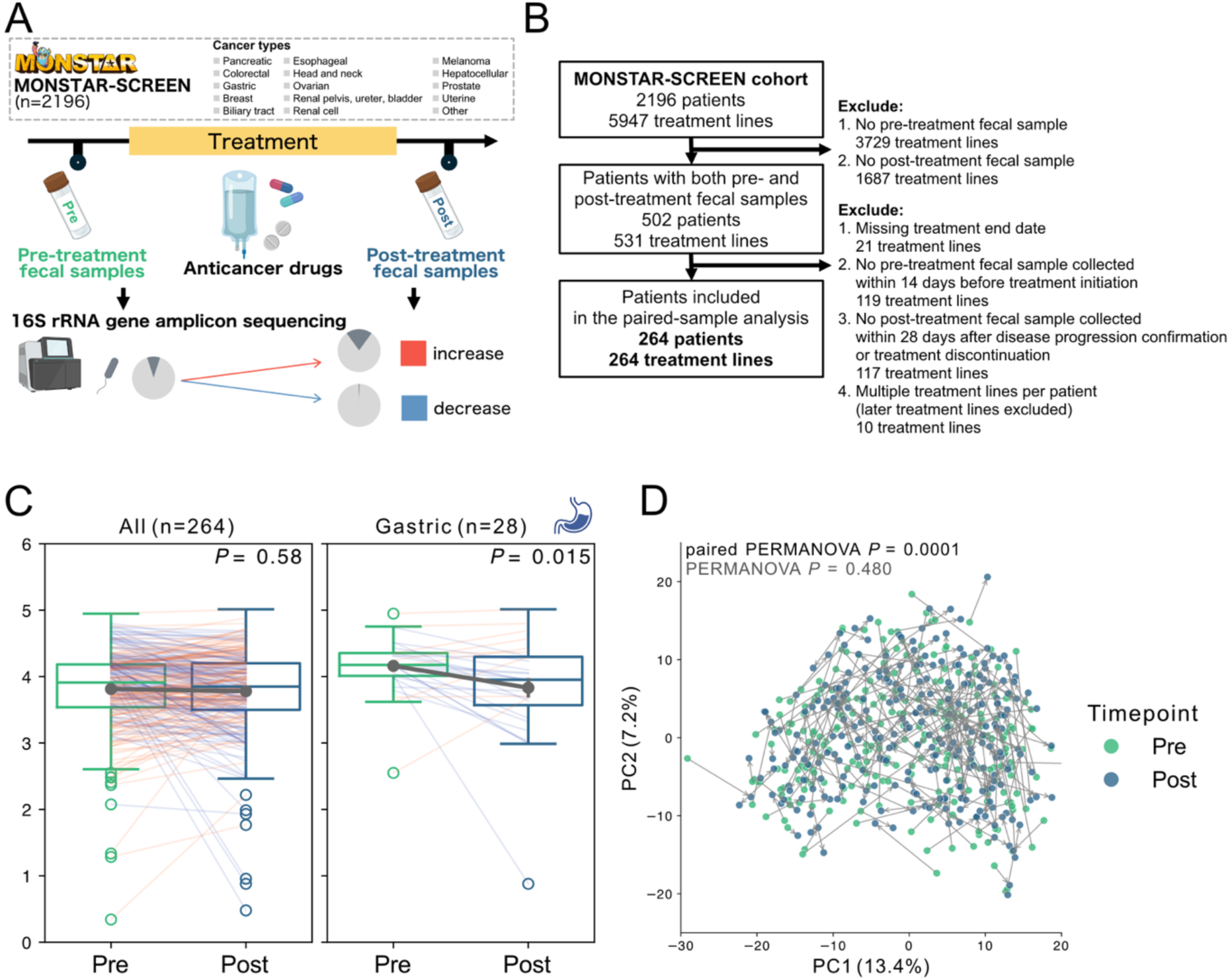
Longitudinal shifts in gut microbiome diversity and composition across the cohort. (A) Study overview (created with BioRender.com). (B) CONSORT diagram. A total of 264 patients with paired pre- and post-treatment fecal samples were included (528 samples in total). Pre-treatment fecal samples were collected within 14 days before or on the day of treatment initiation, and post-treatment fecal samples were collected from the date of confirmed disease progression or treatment discontinuation through 28 days thereafter. (C) Boxplots of Shannon diversity in paired samples (left) and gastric cancer (right). *P* values were calculated using the paired Wilcoxon signed-rank test. Means and standard errors are indicated. (D) PCoA based on the Aitchison distance. Differences between pre- and post-treatment samples were assessed using unpaired and paired PERMANOVA.

**Table 1.** Patient characteristics.

|  |  | mean [range]; n (%) |
| --- | --- | --- |
| **Age** |  | 64 [24–86] |
| **Sex** | Male | 148 (56 %) |
|  | Female | 116 (44 %) |
| **BMI** |  | 22 [12.7–38.8] |
| **Presence of primary tumor** | Yes | 148 (56 %) |
|  | No | 107 (41 %) |
|  | Unknown | 9 (3.4 %) |
| **Line of therapy** | 1 | 183 (69 %) |
|  | 2 | 31 (12 %) |
|  | 3 | 30 (11 %) |
|  | ≥ 4 | 20 (8 %) |
| **Cancer type** | Pancreatic cancer | 59 (22 %) |
|  | Colorectal cancer | 50 (19 %) |
|  | Gastric cancer | 28 (11 %) |
|  | Breast cancer | 24 (9 %) |
|  | Biliary tract cancer | 17 (6 %) |
|  | Esophageal cancer | 15 (6 %) |
|  | Head and neck cancer | 14 (5 %) |
|  | Ovarian, fallopian tube, and peritoneal cancers | 13 (5 %) |
|  | Cancer of renal pelvis, ureter, and bladder | 8 (3 %) |
|  | Renal cell carcinoma | 7 (3 %) |
|  | Malignant melanoma | 7 (3 %) |
|  | Hepatocellular carcinoma | 7 (3 %) |
|  | Primary neuroendocrine tumors/cancers of the digestive tract | 6 (2 %) |
|  | Prostate cancer | 4 (2 %) |
|  | Uterine cancer | 2 (1 %) |
|  | Other | 3 (1 %) |
| **Total** |  | 264 |

Notably, among the four cancer types with ≥ 20 cases, gastric cancer exhibited a significant reduction in diversity (mean, 4.16 vs. 3.83; *P* = 0.015; Figure 1C, right, and Figure S1B). Because PPI use has been associated with gut microbiome alterations [29] and was more common in upper gastrointestinal cancers in this cohort [24], we examined whether it could explain this decline. However, no significant diversity differences were observed between PPI users and non-users (Figure S1C). Beta diversity analysis using Aitchison distance demonstrated no discrete separation between pre- and post-treatment samples in the PCoA, and unpaired PERMANOVA detected no significant differences (*P* = 0.480). In contrast, after accounting for inter-individual variation, paired PERMANOVA detected a significant longitudinal difference *(P* = 0.0001; Figure 1D). A similar pattern was observed across cancer types with at least 20 cases (Figure S1D), highlighting the importance of within-patient comparisons over simple pre/post contrasts. Notably, the distribution of within-patient Aitchison distances was well described by a two-component Gaussian mixture model in the full cohort (Figure S1E), whereas larger within-patient distances were not enriched in any cancer type with ≥ 20 cases (Figure S1F). Together, these results suggest that the observed heterogeneity was not primarily driven by cancer type and may instead reflect patient-specific microbiota alterations.

### Identification of microbial clusters associated with treatment and clinical characteristics

To dissect the drivers of this heterogeneity, we clustered the patients based on genus-level compositional alterations (Figure 2A). K-means clustering classified the patients into six groups (Figure 2B). The magnitude of microbiota compositional shifts—quantified using the Aitchison distance—differed significantly among clusters, with Cluster 6 demonstrating the largest shifts and Cluster 2 the smallest (*P* = 2.7 × 10^-21^; Figure 2C). Differential abundance analysis using ALDEx2 [30] identified the cluster-specific taxonomic signatures. No significant taxonomic alterations were detected in Cluster 2, whereas Cluster 5 demonstrated increases in Ruminococcaceae, *[Eubacterium] coprostanoligenes* group, and *Butyricicoccus*, together with a decrease in *Enterococcus*. In contrast, Cluster 6 demonstrated an increase in *Enterococcus* and decreases in *Bacteroides*, *Oscillibacter*, and *Blautia* (BH-adjusted *P* < 0.05, ALDEx2; Figure 2D).

**Figure 2.**
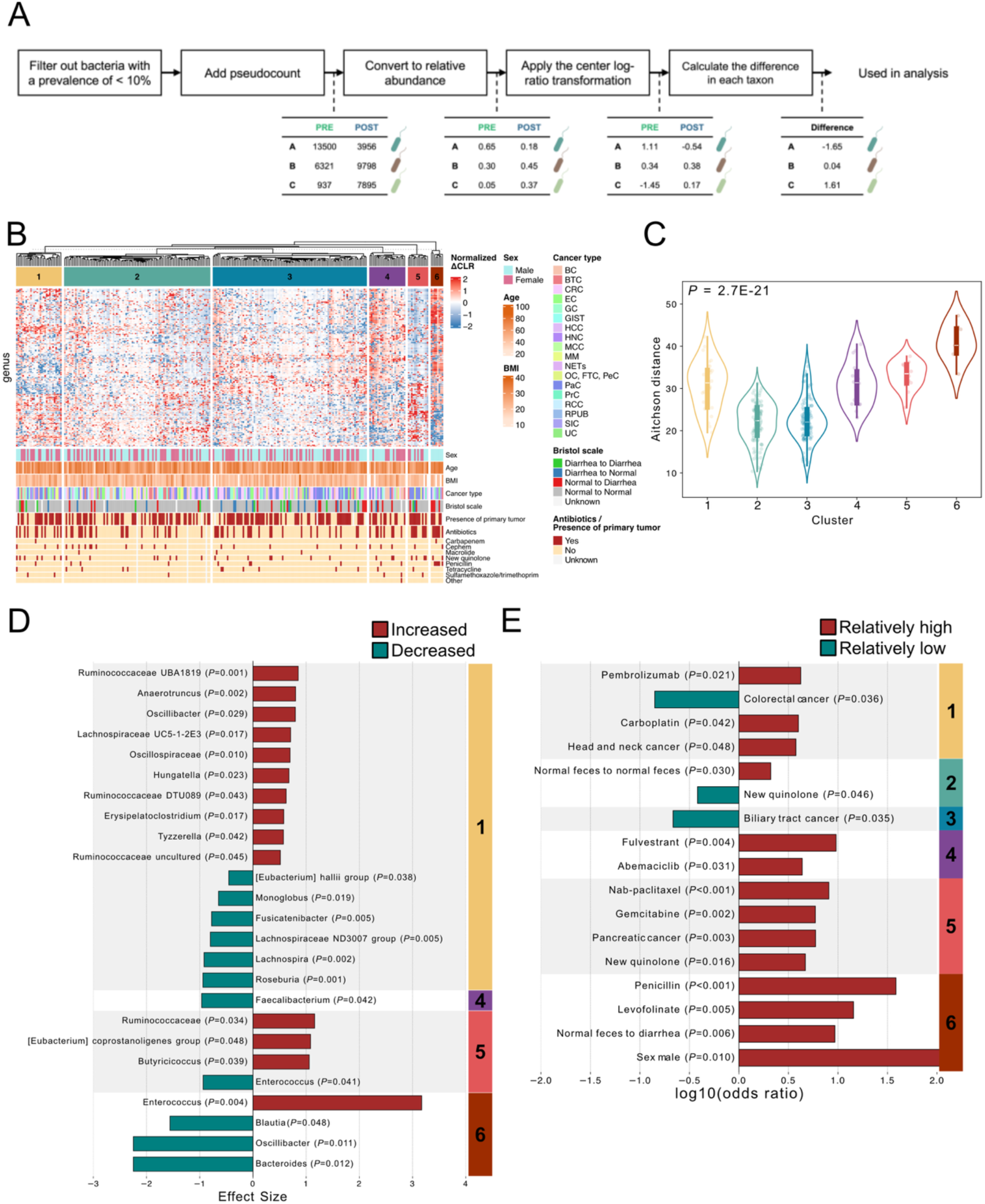
Patient clusters with microbiota compositional alterations. (A) Workflow for calculating genus-level compositional alterations between paired pre- and post-treatment samples. (B) Heatmap of genus-level compositional alterations using k-means clustering, identifying six patient clusters. The clinical characteristics, including sex, age, body mass index, cancer type, Bristol stool scale category, presence of primary tumor, and antibiotic exposure, are annotated below. (C) Violin plots of within-patient Aitchison distances across the clusters. *P* values were calculated using the Kruskal–Wallis test. (D) Cluster-specific taxonomic alterations identified using paired ALDEx2 analysis. Only genera with Benjamini–Hochberg (BH)-adjusted P values < 0.05 are shown, and displayed *P* values are BH-adjusted. (E) Clinical enrichment analysis comparing each cluster with all the remaining clusters combined. Only categorical variables with *P* < 0.05 are presented. BC, breast cancer; BTC, biliary tract cancer; CRC, colorectal cancer; EC, esophageal cancer; GC, gastric cancer; GIST, gastrointestinal stromal tumor; HCC, hepatocellular carcinoma; HNC, head and neck cancer; MCC, Merkel cell carcinoma; MM, malignant melanoma; NETs, neuroendocrine tumors/cancers of the digestive tract; OC/FTC/PeC, ovarian cancer/fallopian tube cancer/peritoneal cancer; PaC, pancreatic cancer; PrC, prostate cancer; RCC, renal cell carcinoma; RPUB, renal pelvis, ureter, and bladder cancer; SIC, small intestine cancer; and UC, uterine cancer.

Clinical enrichment analysis revealed that these clusters were associated with distinct clinical characteristics (Figure 2E; Table S3). Cluster 5 was associated with pancreatic cancer, gemcitabine plus nab-paclitaxel, and new quinolone use, whereas Cluster 6 was enriched for penicillin exposure, transition from normal stool to diarrhea, levofolinate use, and male sex (*P* < 0.05, Figure 2E), and was associated with older age (*P* < 0.05, Table S3). These findings indicate that larger microbiota compositional shifts are associated with treatment-related exposures, including antibiotics and anticancer regimens, and gastrointestinal alterations, such as the transition from normal stool to diarrhea.

### Drug–bacteria association patterns reflect shared mechanisms of action

To estimate associations between anticancer drugs and microbiota alterations while adjusting for clinical covariates, we used multivariable linear regression models incorporating 22 drugs and relevant clinical variables. Multicollinearity was limited, with a maximum VIF of 6.14 (Table S4). This analysis generated a catalog of associations between anticancer drugs and bacterial genera, summarized as a regression coefficient matrix (Figure 3A; Table S5). The catalog captured known associations, including penicillin-associated increases in *Enterococcus* [33, 34] and diarrhea-associated increases in *Veillonella* and *Sutterella* [35] (Figure S2A).

**Figure 3.**
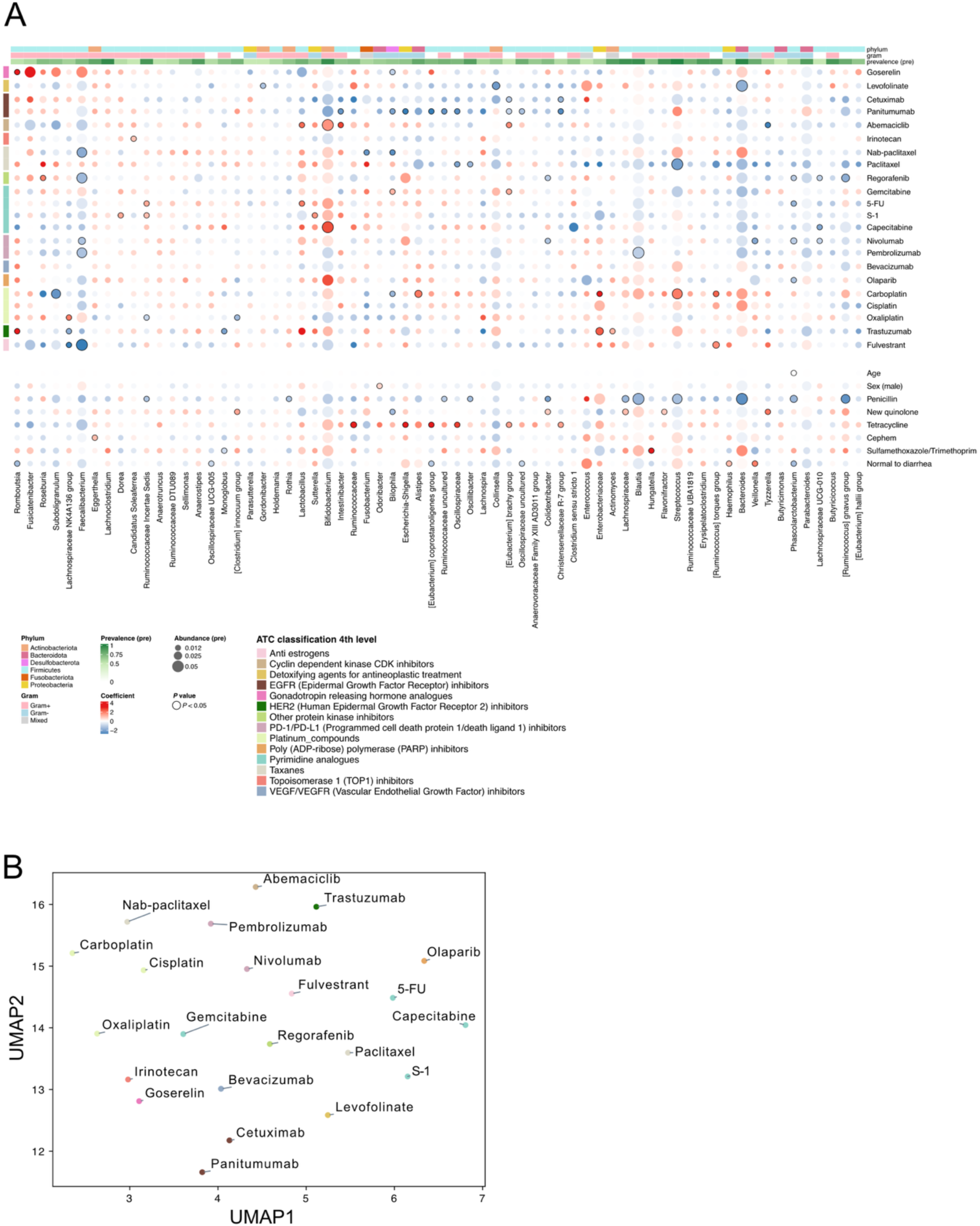
Catalog of drug–bacteria associations across anticancer drugs. (A) Regression coefficient matrix summarizing the associations between anticancer drugs and genus-level microbiota alterations. Coefficients were estimated using separate multivariable linear regression models for each genus, with within-patient change in the centered log-ratio-transformed abundance as the outcome. Anticancer drugs are annotated using Anatomical Therapeutic Chemical (ATC) Classification, 4th level [31]. Bacterial taxa are annotated by phylum, Gram-stain category, as defined in the JGI GOLD database [32], and pre-treatment prevalence. The circle size indicates the mean relative abundance in pre-treatment samples among patients exposed to each anticancer drug. Circles denote associations with a nominal *P* < 0.05 in the regression models. (B) UMAP projection of drug-level association profiles derived from the regression coefficient matrix. The colors indicate ATC Classification, 4th level.

A UMAP projection of these associations demonstrated that drugs with shared mechanisms of action were associated with similar microbiota change profiles (Figure 3B). Specifically, EGFR inhibitors (panitumumab and cetuximab) clustered closely, and a similar pattern was observed for ICIs (nivolumab and pembrolizumab), suggesting class-level similarity in drug–bacteria association patterns.

### Targeted analysis of EGFR inhibitors and ICIs

Focusing on these clinically important drug classes, regression analysis revealed that EGFR inhibitors (panitumumab and cetuximab) were commonly associated with reductions in the Christensenellaceae R-7 group (*P* = 0.002 and 0.034, respectively), Oscillospiraceae (*P* = 0.025 and 0.229, respectively), Ruminococcaceae uncultured (*P* = 0.006 and 0.098, respectively), and *Intestinibacter* (*P* = 0.05 and 0.154, respectively; Figure 4A). In the exposed patients (n = 17), paired analysis of rarefied read counts supported these decreasing trends, although they did not reach statistical significance (Figure 4B, Figure S3A). In contrast, regression analysis associated ICI therapy (nivolumab and pembrolizumab) with a shared trend toward *Lactobacillus* enrichment (*P* = 0.316 and 0.182, respectively) and *Faecalibacterium* depletion (*P* = 0.046 and 0.019, respectively; Figure 4C). In the exposed patients (n = 49), paired analysis of rarefied read counts confirmed a significant reduction in *Faecalibacterium* (*P* = 0.006), whereas the increase in *Lactobacillus* did not reach statistical significance (Figure 4D). These results indicate shared patterns of microbiota alterations within the EGFR inhibitor and ICI classes.

**Figure 4.**
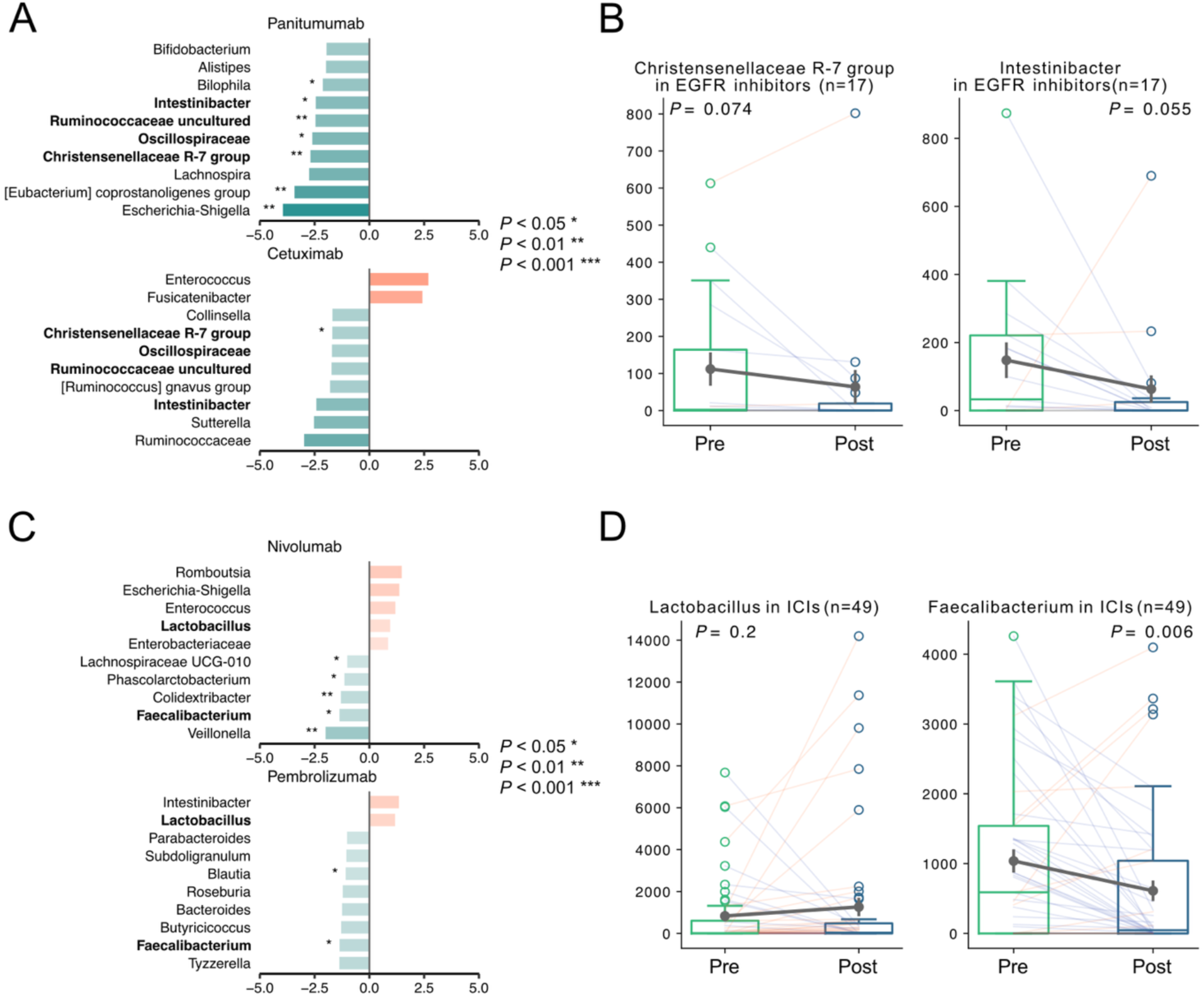
Microbiota alterations associated with EGFR inhibitors and ICIs. (A) Top 10 genera associated with EGFR inhibitors (panitumumab and cetuximab), ranked by the absolute regression coefficient. *P* values were obtained from the regression models. (B) Boxplots of rarefied read counts before and after treatment for the Christensenellaceae R-7 group (left) and *Intestinibacter* (right) in patients exposed to EGFR inhibitors (n = 17). *P* values were calculated using the paired Wilcoxon signed-rank test. Means and standard errors are indicated. (C) Top 10 genera associated with ICIs (nivolumab and pembrolizumab), ranked by the absolute regression coefficient. *P* values were obtained from the regression models. (D) Boxplots of rarefied read counts before and after treatment for *Lactobacillus* (left) and *Faecalibacterium (*right) in patients exposed to ICIs (n = 49). *P* values were calculated using the paired Wilcoxon signed-rank test. Means and standard errors are indicated.

## Discussion

In this study, we analyzed 528 paired fecal samples from 264 patients in the MONSTAR-SCREEN cohort to characterize gut microbiota alterations across diverse cancer types and anticancer therapies. Although the influence of the gut microbiome on cancer treatment response has been increasingly reported [2–8], most longitudinal cohorts have remained limited to specific cancer types or individual regimens [17–20]. To our knowledge, this study represents one of the largest cross-cancer longitudinal analyses to date, using a unified analytical pipeline, enabling systematic comparison of microbiota alterations across multiple therapeutic classes.

At the cohort level, pre- and post-treatment fecal microbiome profiles did not form a simple dichotomy (Figure 1), but longitudinal within-patient alterations varied among individuals. Patients exhibiting the most notable compositional shifts were significantly enriched for antibiotic exposure, diarrhea, and specific treatment regimens (Figure 2). Notably, the increase in *Enterococcus* associated with penicillin-class antibiotics—primarily piperacillin sodium/tazobactam—was consistent with established antibiotic-associated microbiome perturbations [33, 34], supporting the biological plausibility of our analytical approach.

Expanding on these observations, we constructed a multivariable regression-based catalog of drug–bacteria associations adjusted for available clinical confounders (Figure 3). This systematic reference suggested that drugs with shared mechanisms of action were associated with concordant microbiota change profiles. For example, EGFR inhibitors, including panitumumab and cetuximab [36], and ICIs, including nivolumab and pembrolizumab [37], exhibited similar microbial signatures, indicating that the shared pharmacological mechanism may contribute to similar selective pressures on the gut ecosystem.

For EGFR inhibitors, we identified concordant reductions in the Christensenellaceae R-7 group, Oscillospiraceae, Ruminococcaceae uncultured, and *Intestinibacter* (Figure 4 and Figure S3). The Christensenellaceae R-7 group has been associated with favorable metabolic profiles and healthy aging phenotypes [38]. Given that EGFR signaling contributes to intestinal epithelial homeostasis and barrier integrity [39], pharmacological inhibition of EGFR may alter the intestinal mucosal environment. Because the EGFR inhibitors analyzed here target the host EGFR protein rather than bacterial components [36], the observed microbiota alterations are more likely to arise, at least in part, from host-mediated effects rather than direct antibacterial activity. Therefore, we hypothesize that EGFR blockade-associated changes in the intestinal environment may contribute to the depletion of commensal taxa, including the Christensenellaceae R-7 group.

For ICI therapy, we observed a significant reduction in *Faecalibacterium*, together with a trend toward *Lactobacillus* enrichment (Figure 4). *Faecalibacterium*, specifically *F. prausnitzii*, has been repeatedly associated with favorable ICI outcomes and has experimental support as a potential enhancer of ICI-mediated antitumor immunity [3, 40–42]. Because most samples in this subset were collected during disease progression (48/49, 98.0 %), these profiles may reflect a progression-associated microbiome state rather than the direct effect of the drug itself. This aligns with previous longitudinal evidence showing decreasing trends in Lachnospiraceae and *Faecalibacterium* during the development of secondary resistance in patients with advanced thoracic cancers receiving anti-PD-1 therapy [19]. In summary, these data highlight the necessity of longitudinal microbiome monitoring to identify clinically relevant windows for microbiome-directed interventions.

This study has several limitations. First, it remains challenging to disentangle whether microbiome shifts are driven by direct effects of anticancer drugs on bacterial growth or metabolism, or by indirect effects through disease progression and host-mediated factors. This is particularly relevant for regimens used preferentially in specific malignancies, where confounding by indication is inherent. Second, post-treatment sampling time—often coinciding with clinical progression—may bias observations toward a resistant phenotype. Third, certain unmeasured confounders cannot be fully adjusted. Finally, because microbiome data are inherently compositional, future studies incorporating absolute quantification (microbial load) are required to enhance the precision of these interpretations [43–45].

## Conclusions

This study provides a catalog of microbiota alterations associated with anticancer therapies across multiple therapeutic classes. Using a unified analytical framework, we identified therapy-associated shifts in the gut ecosystem, including patterns shared by drugs with similar mechanisms of action. These findings provide a resource for future mechanistic studies and may support the timing and application of microbiome-targeted strategies in cancer therapy.

## Declarations

### Ethics approval and consent to participate

The study adhered to the Declaration of Helsinki and Japanese Ethical Guidelines for Medical and Health Research Involving Human Subjects. The protocol was approved by the institutional review board of each participating institution and registered with the University Hospital Medical Information Network Clinical Trials Registry (UMIN000036749). All participants provided written informed consent.

### Consent for publication

Not applicable.

### Availability of data and materials

Raw sequencing data and associated sample metadata are available in the DDBJ Sequence Read Archive under the BioProject accession number PRJDB39670. Additional datasets supporting the conclusions of this study are included in this article and its additional files. The code used in this study is available at https://github.com/kyokokurihara/anticancer-drug-gut-bacteria-catalog.

## Authors’ contributions

**K.K.**: Methodology, Data Curation, Formal Analysis, Visualization, and Writing - Original Draft. **S.A.S.**: Methodology, Formal Analysis, and Writing - Review & Editing. **K.S.**: Writing - Review & Editing. **N.I.**: Data Curation. **S.H.**: Data Curation. **T.F.**: Project Administration and Writing - Review & Editing. **Y.N.**: Project Administration. **S.K.**: Writing - Review & Editing. **H.B.**: Project Administration. **T.Y.**: Funding Acquisition and Project Administration. **K.T.**: Writing - Review & Editing. **R.Y.**: Project Administration, Conceptualization, and Writing - Original Draft. All authors reviewed the manuscript.

## Acknowledgements

We thank all patients and their families for participating in this study. We also thank the clinicians, clinical research nurses, clinical research coordinators, and other research personnel at each site for their contributions, and the members of the Translational Research Support Section at the National Cancer Center Hospital East. Computational analyses were performed on the KASHIWARP server at the National Cancer Center. We thank Naoki Amano for his valuable feedback and Editage (https://www.editage.jp/) for English language editing.

## List of abbreviations

anti–PD-1: anti–programmed cell death protein 1
ASV: Amplicon sequence variant
BMI: Body mass index
CLR: Centered log-ratio
EGFR: Epidermal growth factor receptor
ICI: Immune checkpoint inhibitor
PCoA: Principal coordinates analysis
PCR: Polymerase chain reaction
PERMANOVA: Permutational multivariate analysis of variance
PPI: Proton pump inhibitor
rRNA: Ribosomal RNA
UMAP: Uniform manifold approximation and projection
VIF: Variance inflation factor

---

**Figure S1.**
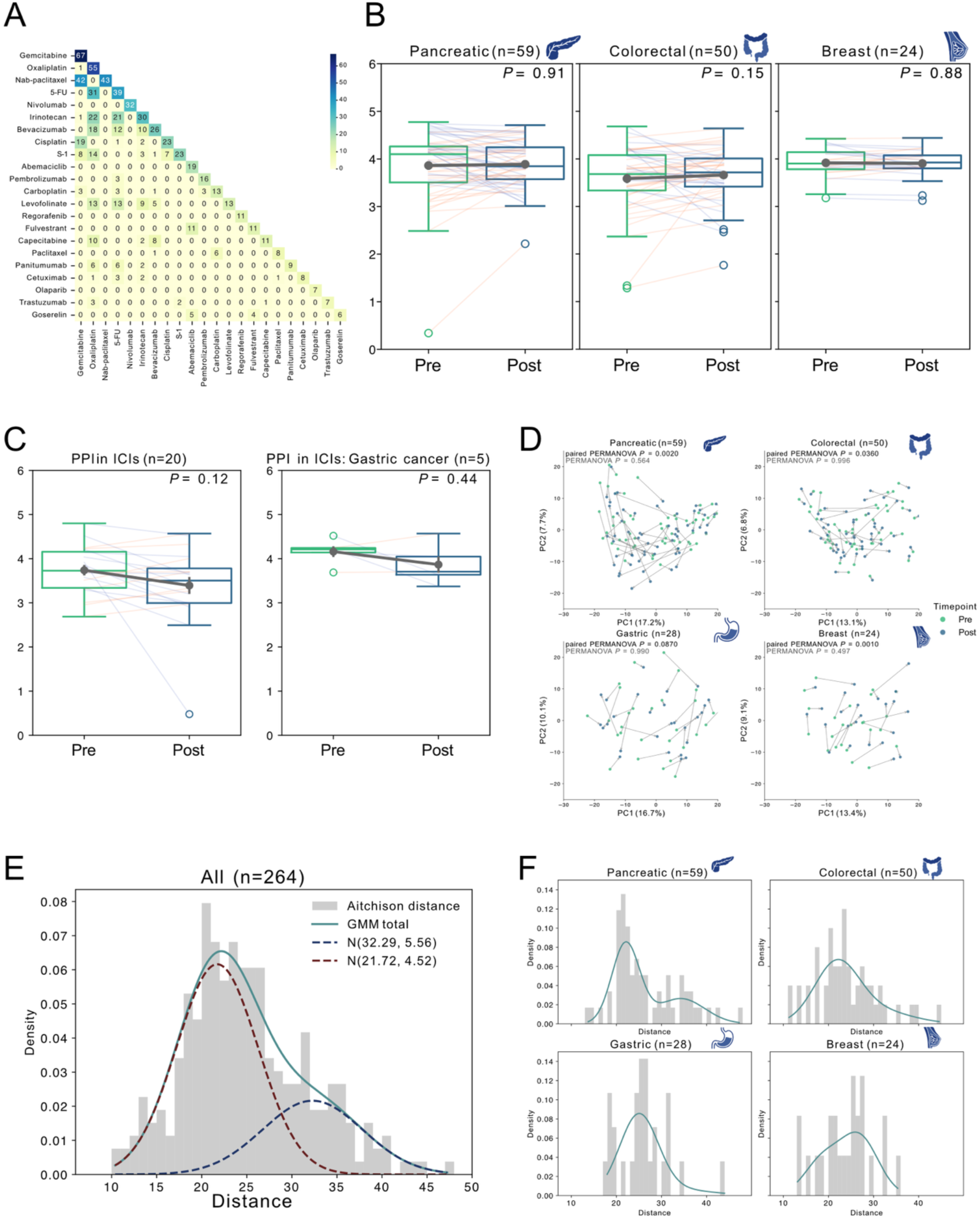
Supplementary analyses of longitudinal diversity alterations. (A) Heatmap of co-administered anticancer drugs used in at least five patients. (B) Boxplots of Shannon diversity in paired pre- and post-treatment samples from pancreatic cancer (left), colorectal cancer (middle), and breast cancer (right). *P* values were calculated using the paired Wilcoxon signed-rank test. Means and standard errors are indicated. (C) Boxplots of Shannon diversity in immune checkpoint inhibitor-treated patients with proton pump inhibitor exposure (left) and in the gastric cancer subset (right). *P* values were calculated using the paired Wilcoxon signed-rank test. Means and standard errors are indicated. (D) PCoA based on Aitchison distance, stratified by cancer type, with at least 20 cases. Differences between pre- and post-treatment samples were assessed using unpaired and paired PERMANOVA. (E) Distribution of within-patient Aitchison distances across the cohort fitted with a two-component Gaussian mixture model. (F) Distribution of within-patient Aitchison distances for cancer types with at least 20 cases.

**Figure S2.**
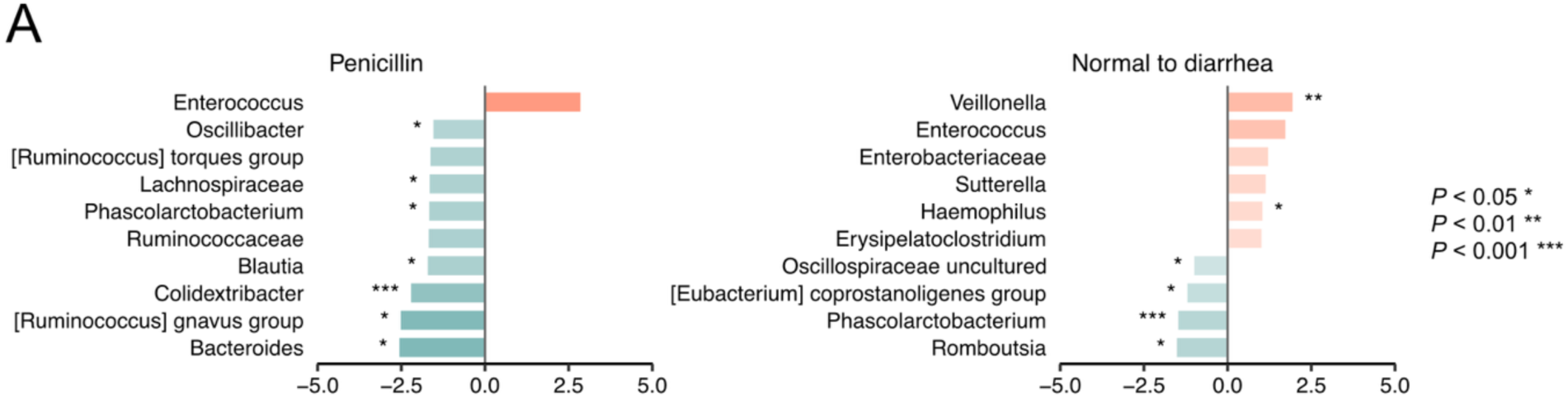
Antibiotic- and diarrhea-associated microbiota alterations captured using regression-based catalog. (A) Top 10 genera associated with penicillin exposure (left) and transition from normal stool to diarrhea (right), ranked by absolute regression coefficient. *P* values were obtained from the regression models.

**Figure S3.**
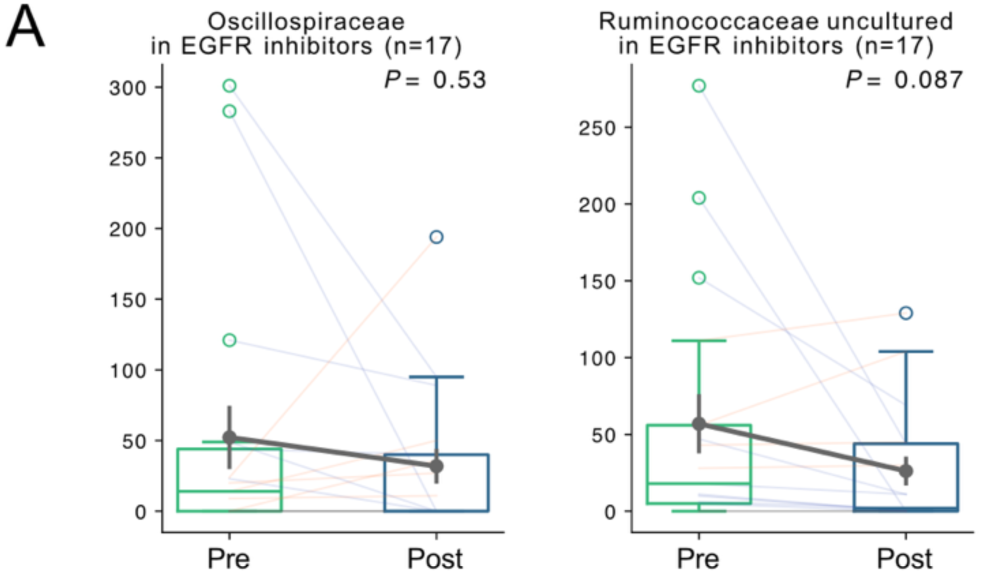
Additional taxa associated with EGFR inhibitor exposure. (A) Boxplots of rarefied read counts before and after treatment for Oscillospiraceae (left) and Ruminococcaceae uncultured (right) in patients exposed to EGFR inhibitors (n = 17). *P* values were calculated using the paired Wilcoxon signed-rank test. Means and standard errors are indicated.

## Additional files

**Table S1. Detailed breakdown of cancer types and treatment regimens in the analysis cohort.**

Cancer type and treatment regimen combinations in the paired-sample analysis cohort are represented.

File format: .xlsx

**Table S2. Classification of antibiotic exposure.**

Antibiotic exposure is grouped into eight classes: cephem, new quinolone, penicillin, tetracycline, carbapenem, macrolide, sulfamethoxazole/trimethoprim, and others.

File format: .xlsx

**Table S3. Associations between patient clusters and clinical characteristics.**

Association analyses between k-means-defined patient clusters and clinical characteristics. Each cluster was compared with all the remaining clusters combined. Continuous variables were analyzed using the Mann–Whitney U test, and categorical variables were analyzed using Fisher’s exact test. Categories represented by fewer than five patients were excluded.

File format: .xlsx

**Table S4. VIF for the variables included in the multivariable regression models.**

VIF for the anticancer drugs and clinical covariates included in the multivariable regression models. File format: .xlsx

**Table S5. Regression coefficients, nominal *P* values, and pre-treatment prevalence of drug–bacteria associations.**

Regression coefficients and nominal *P* values from separate multivariable linear regression models for each genus, together with the pre-treatment prevalence. Genera with a pre-treatment prevalence of at least 50 % were used for the primary analysis.

File format: .xlsx

